# Biodegradation of polyester polyurethane by *Aspergillus flavus* G10

**DOI:** 10.1101/2020.06.25.170654

**Authors:** Sehroon Khan, Sadia Nadir, Yang Dong, Douglas A. Schaefer, Peter E. Mortimer, Heng Gui, Afsar Khan, Mingming Yu, Shahid Iqbal, Jun Sheng, Jianchu Xu

## Abstract

Polyurethanes (PU) are integral to many aspects of our daily lives. Due to the extensive use of and difficulties in recycling or reusing PU, it mostly accumulates as waste. Various bacteria and fungi have been reported to degrade PU. We examined the fungus *Aspergillus flavus G10* in that regard, after isolating it from the guts of *Gryllus bimaculatus*, a common cricket species. We observed surficial and chemical changes of PU with atomic force microscopy, scanning electron microscopy, and attenuated total reflectance Fourier-transform infrared spectroscopy. We measured physical changes as loss in tensile stress, stretching force, and weight of PU after incubations. Fungal hydrolysis of urethane bonds in the polymer backbone was demonstrated by detecting the formation of methylene di-aniline during incubations. Trapped CO_2_ during incubations equaled 52.6% of PU carbon. Biodegradation of PU was maximal by fungi cultured on a malt extract medium at 25 °C, pH 12, and 14:10 hrs light to dark ratio. Pretreating PU films with UV light or 1% FeSO_4_ or NaCl solutions further enhanced the rate of biodegradation. A range of techniques are needed to fully characterize the degradation of PU or other plastic polymers and to optimize conditions for their microbial degradation.

## Introduction

Annual production of plastics increased to 355 million tones (Mt) in 2017 due to societal and commercial benefits [1]. 9150 Mt of primary plastics have been produced from 1950 to 2015, resulting in 6945 Mt of plastic waste on the surface of earth [2]. However, not all plastic waste products are landfilled, incinerated, or recycled. Environmental persistence of plastic waste can produce pollutants that have been shown to be threatening to life in different ecosystems [3, 4, 5, 6].

Polyurethane (PU) is a wisely used plastic, accounting for about 7% of total plastic production, with a global annual production of 12 Mt [2]. In China alone the annual production of polyurethane reached 7.5 Mt in 2011 [7]. Incineration and chemical recycling of PU waste are not feasible solutions to PU recycling. In light of this, microbial degradation is one of the most environmentally friendly and cost-effective ways to manage PU waste [7].

Many studies examine the microbial degradation of plastics [8, 9, 10] or PU biodegradation alone [11, 12, 13, 14, 15, 7]. Matsumiya et al. [16] reported a 27.5% weight decrease in ether-type PU after 10 weeks’ incubation with the fungus *Alternaria sp*.

Various methods are used to visualize types and extent of polyurethane biodegradation by fungi or other microbes. According to Khan et al. [11], the holes, pits, and other signs of erosion that were revealed in the scanning electron microscopic (SEM) analysis of the PU films showed biodegradation. Bombelli et al. [17] used Atomic Force Microscopy (AFM) analysis to analyze the surface degradation of *Galleria mellonella* homogenate on the surface of PE. Others used Attenuated Total Reflection Fourier Transform Infrared (ATR-FTIR) spectra to determine chemical alterations in treated and untreated plastics samples [11, 18, 19, 20, 21]. Loss of tensile strength also indicates biodegradation [22, 23, 24].

Khan et al. [11] used tween 20 and 80 to reduce the hydrophobicity of the PU film surface to make it accessible for fungal spore attachment. Khan et al. [11] demonstrated high enzyme activity at 37°C in acidic liquid broth culture of *A. tubingensis*. Idnurm and Heitman [25] demonstrated that the endophytic fungus was able to sense darkness and use that as a signal to induce virulence, suggesting the possibility that exposure to different light-to-dark ratios could affect biodegradation.

Methylene di-aniline (MDA) is known to be the enzymatic biodegraded compound of polyurethanes used as a model compound to analyze the degradation of polyurethanes by enzymes [26]. The fungal strain PURDK2 was found to be able to hydrolyze both urethane and urea bonds in ether-type PU and utilize degraded compounds as a carbon source [16].

Next, we isolated the cricket gut fungus *A. flavus* G10 and tested its biodegradation with culture plate and liquid broth methods. Biodegradation of PU was observed using atomic force microscopy, scanning electron microscopy, and attenuated total reflectance Fourier-transform infrared spectroscopy. We also optimized biodegradation by pretreating PU film with UV exposure and incubation in dilute solutions of NaCl, MgSO_4_, ZnSO_4_, FeSO_4_, and CuSO_4_ salts to determine optimal PU biodegradability. Furthermore, production of CO_2_ and MDA were determined. Our results bring novel insights into the field of biodegradation and offer practical significance for environmental remediation.

## Materials and methods

We used PU beads, with the chemical formula “poly [4, 4’-methylenebis (phenylisocyanate)-alt-1, 4-butanediol/di (propylene glycol)/polycaprol-actone],” (Aldrich Chemical Company, Inc. USA). The PU beads (1g) were dissolved in tetrahydrofuran (THF) (Sigma-Aldrich) by being shook in a conical flask for 12 hours at 150 rpm at n room temperature. Dissolved PU was poured into glass Petri dishes for solvent evaporation to make PU films. Two types of PU films were prepared by two different drying methods. Dissolved PU air dried at 50% humidity made foamy films, and when dried in a desiccator, PU films formed were transparent [11].

### Biodegradation test on solid medium

Solid medium biodegradation was performed for PU films following the method of Khan et al. [11]. Both foamy and transparent types PU films were used for biodegradation analysis. Six MEA plates were prepared according to the manufacturer’s instructions. Three MEA plates at pH 7.6, containing transparent PU or foamy PU films, were inoculated with 0.4 ml spore suspension (1×10^5^ spores/ml) of *A. flavus* G10. Those plates were incubated at 30 °C for 28 days. MEA plates containing PU films with no fungal inoculum were used as controls.

### Biodegradation in liquid medium

*A. flavus* G10 was tested for PU biodegradation in a broth culture of malt extract at pH 7.6. Three Erlenmeyer flasks containing 300 ml of malt extract broth were inoculated with 2 ml/flask spore suspension (1× 10^5^ spores/ml) of *A. flavus* G10. Two PU transparent films (90 mm) were cut longitudinally, sterilized with UV radiation for 5 minutes by exposing the PU films inside laminar flow hood, and were added into each of three Erlenmeyer flasks and incubated at 30 °C (150 rpm) using a Zhicheng incubator shaker (ZWF 200). After 24, 48, and 72 hours of incubation, samples were collected and brushed gently in sterilized water for one minute. Three PU films immersed in malt extract (ME) broth at pH 7.6, having no fungal inocula, were controls. The liquid biodegradation test was performed for transparent PU films following the method of Khan et al. [11] and repeated for foamy PU films in triplicates. To visualize biodegradation, scanning electron microscopy (SEM) studies were carried out by using the ZEISS Sigma 300 with accelerating voltage from 3 to 7KV; magnification from 50 X to 4500 X; and resolution from 200 μm to 200 nm for both transparent and foamy PU films.

### Biodegradation rates of PU film

75 transparent PU films were prepared and exposed to UV for 10 minutes in a laminar flow hood, with 5 minutes on each surface. The initial weight of each film was then measured, and the films were next placed on the *A. flavus* G10 culture in covered Petri dishes. For 15 consecutive weeks, five PU films were removed and washed gently with sterilized water to remove mycelial mass from PU film surfaces. PU films were then dried at room temperature and reweighed. Using the mean values of the 15 weeks, the % DE of the fungus was determined according to the following formula:

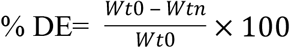

*Wt*_*0*_ is the initial weight of PU film and *Wtn* is the weight of PU film after each incubation period (n =1 to 15 weeks). Mean % DE values accumulated over time were treated as efficiency of fungal degradation.

### Surface topological studies of transparent PU films

To assess the effects of *A. flavus* G10 on the surface topology of transparent PU films, mycelia of *A. flavus* G10 were crushed with a mortar and pestle, spread onto three transparent PU films, and then incubated at 37 °C for two, four, or six hours. PU films were gently brushed with sterilized water and stored at room temperature. Three PU films were established as controls by spreading autoclaved crushed hyphal masses on PU films. Atomic force microscopy (AFM) analysis was used to visualize the surface topology of PU films using AFM (Dimension Icon, Veeco, USA). The AFM was set at 2000 nm and three-dimensional images were scanned.

### Chemical analysis of the biodegradation of the PU films

To explore bond breakage or formation, we performed infrared spectral analysis following the method of Khan et al. [11]. Three replicates of incubated and control transparent PU films were used for analysis. Infrared spectra of these were obtained with an attenuated total reflectance (ATR) accessory for Fourier-transform infrared (FTIR) spectrophotometer (Thermo Scientific Nicolet 10). The spectral range was 4000 to 600 cm^−1^ with a resolution of 4 cm^−1^. Samples were placed on the ATR spot of the additional ATR portion and pressed slowly.

### Mechanical properties of the PU film exposed to *A. flavus* G10

To characterize mechanical properties after biodegradation, transparent PU films (n=11) were exposed to *A. flavus* G10 on a solid medium at 30 °C for a period of four weeks. After every week, three PU films were randomly selected and brushed with sterilized water and stored at room temperature until analysis. This process was continued until the end of 4^th^ week. A set of three transparent PU films that was not exposed to *A. flavus* G10 were controls.

Tensile properties were measured with Biotester 5000 (CellScale, Canada). The amount of force required to stretch PU films (1 cm^2^) longitudinally (gage length L_0_) to double its original length (2L_0_) was calculated for both experimental and control PU films. The Biotester 5000 was set at 0.1 mm/s using displacement control with five repetitions for each, at a temperature of 18±2 °C. Photographs were taken at L_0_ and 2L_0_. From the data obtained from Biotester 5000, other mechanical properties such as tensile stress and strain were calculated as below:

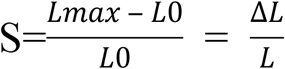

where S is the strain, and L_max_ is the stretched length of the PU film; and

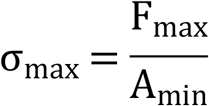

where σ_max_ is the tensile stress, F_max_ is the maximum amount force on the PU film, and A_min_ is the area of the PU film exposed to applied maximum force.

An electronic caliper (Chuanlu Measuring Tools, Co. LTD, Shanghai) was used to measure thickness of the PU films. Measurements were made every week for a period of four weeks (Wk1 to Wk4).

### LC/MS analysis of the PU degraded compounds

Haugen et al. [27] used cholesterol esterase for enzymatic degradation of polyurethane to test the biostability of the polymer and found MDA as a biodegradation product. We determined MDA after the biodegradation of PU with *A. flavus* G10 following the method of Johnson et al. [28]. The mycelium of *A. flavus* G10 along with the degraded PU film residues (5 g) obtained from the test culture plate were added into conical flasks with 20 ml of a 1:1 methanol and ethyl acetate solution. The mixture was filtered through filter paper and solvent was evaporated. 10 mg of obtained solid residue was dissolved in 5 ml acetonitrile, having 0.1% acetic acid to make a suspension. The suspension was centrifuged for 10 minutes at 12000 rpm at room temperature. The supernatant was collected and used for liquid chromatography–mass spectrometry (LC-MS) analysis to determine MDA. Another solution was prepared to use as a negative control for LC/MS by adding mycelium and MEA (5 g) obtained from fungal culture plate without PU in the same way as described above. This was used as a negative control. A standard solution of 4, 4’-Methylenedianiline (MDA) was prepared by dissolving 0.05 g in 1 mL of acetonitrile with 1% acetic acid. The mixture was centrifuged at 12000 rpm for 10 minutes and the supernatant collected for LC-MS analysis. A non-degraded PU film was used as a control.

High performance liquid chromatography (HPLC) was performed using the Agilent 1200 Infinity system equipped with a semi-preparative C-18 column (5 μm, 250 mm × 4.6 mm). A gradient method was developed to analyze the degradation products [28]. A 10% to 100% solution of a mobile phase consisting of 0.05 M NH_4_OAc solution in water (solvent A), as well as HPLC-grade acetonitrile (solvent B), were used to flush the column over a period of 38 minutes at a flow rate 1 ml/min. The UV absorbance of the eluate was monitored at 254 nm and the fractions corresponding to each HPLC peak were collected. Product identities were confirmed by Electronspray ionization mass spectrometry (ESI-MS) (Agilent G6230), using a direct insertion probe.

### Mineralization of PU

The mineralization of PU was determined by measuring the production of CO_2_ following the methods of Chinaglia et al. [29]. The conversion of PU carbons to CO_2_ by *A. flavus* G10 was assessed for incubation periods of 3, 6, 9, and 12 days. A 2000 mL glass jar containing five MEA Petri dishes inoculated with *A. flavus* G10 (400 μl spore suspension (1×10^5^ spores/ml) and covered with PU films were placed in an incubator at 30 °C under 8:16 dark to light ratio for 12 days. All the culture plates were inoculated with *A. flavus* G10 for 10 days before use in this experiment. A 500 ml beaker filled with 300 ml of 1 M KOH (CO_2_ trapping solution) was also placed in the jar besides culture plates. The 2000 ml glass jar was sealed. A glass jar containing non-inoculated MEA Petri dishes with a PU film was used as a negative control while a glass jar containing *A. flavus* G10 (concentration of 1×10^5^ spores/ml) cultured on MEA Petri dishes (400 μl spore suspension) was used as a positive control. The incubation time was 3, 6, 9, and 12 days and three replicates were used for each measurement period. The CO_2_ produced was measured by titrating KOH solutions with 0.5 M HCl using a potentiometric titrator (Orion Star T900, Thermo Scientific, USA). The conversion of PU to CO_2_ was estimated by subtracting the average amount of CO_2_ produced in the blank to the amount of CO_2_ produced in the PU-containing jars.

### Statistical analysis

The averages and standard errors were computed using Microsoft Excel 2016. To compare differences among treatments, we conducted analysis of variance (ANOVA) in R language and when differences were significant, we performed a pairwise difference analyses with a post-hoc analysis using the TukeyHSD function in the package “agricolae” [30] and also used the plotting function in the “ggplot2” package to plot figures. For ANOVA, we checked whether the models met ANOVA assumptions, and if they did not, we transformed. The only case in which the normal distribution of errors was not met was for media treatments, and we performed logit transformation on those data [31].

## Results

### *A. flavus* G10 for PU biodegradation

*Aspergillus flavus* G10 colonized and grew on transparent and foamy PU films, (Fig 1). Surficial fungal growth on surfaces was easily observed with the naked eye. The discoloration of the PU films, along with the holes and pits in the films, were clear markers of biodegradation. Both kinds of PU films were digested in the center of PU films by the fungus while holes, splits, and discolorations can be seen throughout PU films (Figs 1B and D). Both transparent and foamy control PU films appeared plain and smooth (Figs 1A and C). SEM results further confirmed that biodegradation had taken place (Figs 1A-D). Fungal spores, mycelium, and hyphae could be seen growing on foamy PU films (Fig 1D). Similarly, the SEM results of transparent and foamy PU films exposed to the fungus showed that the fungus grew across PU film surfaces, with the film broken into pieces. Cracks in transparent PU film, with mycelia inside cracks, are shown in Fig 1 (D).

**Fig 1.**
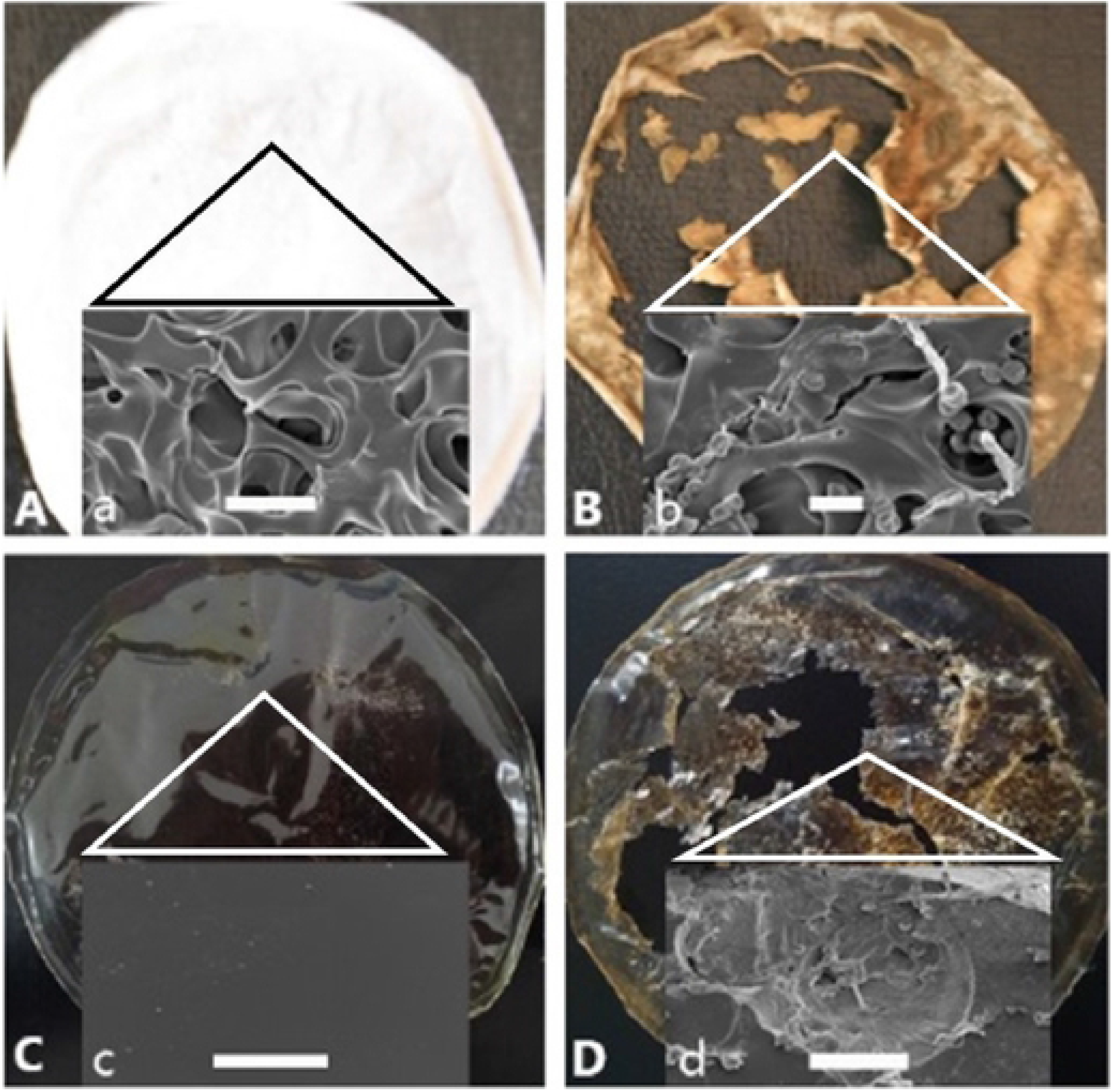
PU biodegradation by *A. flavus G10.* (a) and (c) represent foamy and transparent control PU films respectively that were not exposed to *A. flavus* G10. (b) and (b) represent foamy and transparent PU films, respectively, that were exposed to *A. flavus* G10 for 28 days. The light green/brown color was the growth of fungus on surfaces of PU films. SEM images of foamy PU films (a) Control foamy PU. (b) foamy PU exposed to *A. flavus* G10. Fungal spores, mycelia and hyphae can be seen in the SEM image. (c) Control transparent PU film (d) treated transparent PU films. Scale bars a = 10 μm, b = 4 μm, c = 60 μm, and d= 20 μm.

### Biodegradation efficiency of *A. flavus* G10 by percentage

The percentage weight of the PU films exposed to *A. flavus* G10 decreased over time, from week 1 (Wk 1) to week 15 (Wk 15). Average weight loss of the treated PU films was 1.9% per week (S1 Fig). Biodegradation efficiency (% DE) of PU films exposed to *A. flavus* G10 significantly increased from Wk 1 to Wk 15. (S1 Fig, F1 = 9.368, P = 0.003).

### Surface topology and chemical analysis of PU-degraded films

Atomic Force Microscopy (AFM) images showed that while control transparent PU film surfaces remained smooth (Fig 2A), surfaces of transparent PU films exposed to *Aspergillus flavus* G10 were rough, with deep, wide grooves (Fig 2B-D). The average roughness (Ra) recorded for the PU films exposed to the fungus was 2.66, 2.72, and 6.59 nm after 2, 4, and 6 hours of incubation, respectively. Ra values for control PU film surfaces were 1.13 nm. The cross-sectional view of control PU film (S2 A Fig) was smoother than treated PU films (S2 B-D Fig). The differences (Δ) in the length of the highest peak (Z) and the length of lowest point of the groove ΔZ observed for the treated PU films were approximately 14.68, 17.89, and 44.5 nm after 2, 4, and 6 hours of incubation with the fungus as compared to the ΔZ value of 12.31 nm for the control PU. The surface percentage-bearing ratio for the treated PU films was 38.34, 2.42, and 4.32%, after 2, 4, and 6 hours of incubation respectively, while for control PU films it was 0.13%.

**Fig 2.**
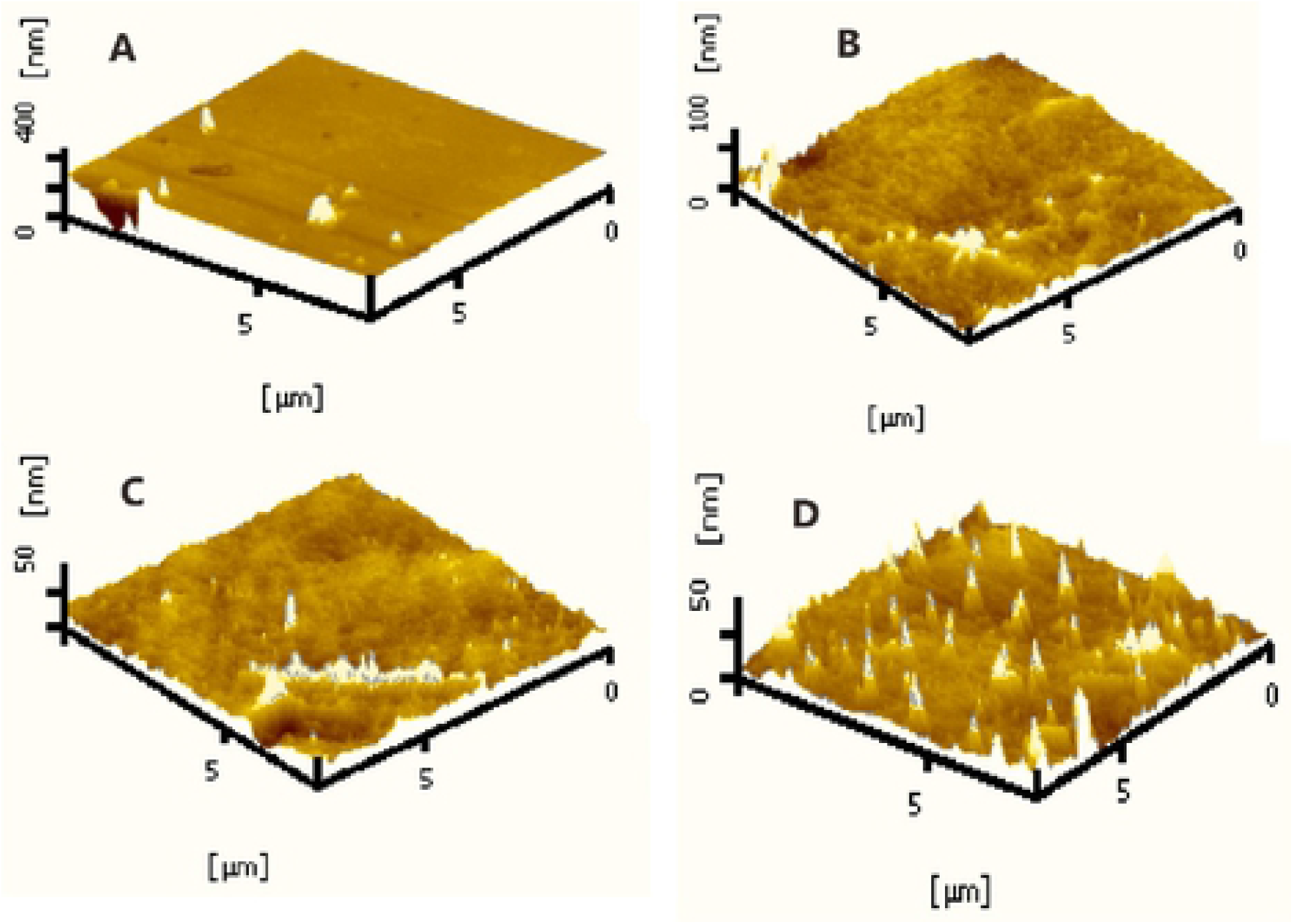
Three-dimensional (3D) topography of the surface of PU films using AFM. (a) 3D surface scan of control PU film. (b-d) 3D surfaces of PU films treated for 2, 4, and 6 hours with mashed *A. flavus.* Scan size is 8nm.

AT-FTIR analysis revealed a notable shift in the spectrum of the foamy PU-treated film (3326.17 cm^−1^) compared to the control (3327 cm^−1^) PU film (S3 A and B Figs), which corresponded with the NH bond deformation. Furthermore, another band, corresponding to CH_2_ asymmetric stretching at 2933.10 cm^−1^ in the control spectra, was observed to have shifted to 2936.64 cm^−1^ in the spectrum of foamy PU film, indicating cleavage of methylene groups on the polymer backbone (S3 A and B Figs). A small band (2866.81 cm^−1^) that appeared on the shoulder of the methylene band corresponding to symmetric CH_2_ stretching had also shifted to 2867.18 cm^−1^ in the test spectrum of foamy PU film. Similarly, a shift in the position of the band representing C-O…H stretching (S3 A and B Figs) was also observed in the spectra of the treated foamy PU film. A sharp band in the control PU film spectrum at 1066.38 cm^−1^ shifted to 1078.17 cm^−1^, representing non-bonded C-O groups. Another prominent change was in the sharpness and intensity of the band at 1726.92 cm^−1^ for the urethane carbonyl group, indicating an increase in the concentration of free urethane carbonyl groups in treated foamy PU films. AT-FTIR analysis of the transparent PU film indicated that the broad band representing the N-H bonding was found at 3322.51 cm^−1^, a shift from the control PU films, which exhibited a band at 3327.93 cm^−1^ in treated PU film (S3 C and D Figs). A second peak in control transparent PU film at 3122 cm^−1^ was found to be decreased in intensity in the treated PU film, indicating NH deformation. The band at 2921.49 cm^−1^, representing CH_2_ asymmetric stretching in the control spectrum, shifted to 2934.91 cm^−1^ (S3 C and D Figs) in treated PU films. Furthermore, a small band representing CH_2_ symmetric stretching at 2866.19 cm^−1^ in control PU films shifted to 2866.43 cm^−1^ in PU films. A sharp band at 1726.82 cm^−1^ representing urethane carbonyl groups in control PU films shifted to 1727.09 cm^−1^ in the treated transparent PU films. Finally, a sharp band in control PU film spectrum at 1066.38 cm^−1^ shifted to 1078.17 cm^−1^ in transparent treated PU films, representing non-bonded C-O groups (S3 C and D Figs).

#### Biodegradation of PU films and its effects on mechanical properties

The stretching force measured on control PU films (0.036 mm thick) was recorded to be 1543.34 mN at Wk0 (S4 A Fig). After one week (Wk1) of incubation with *A. flavus* G10, that decreased to 1403.34 mN. The stretching force decreased further during later incubation, i.e., from Wk2 to Wk4 (S4 Fig), showing significant difference (p= 0.0015) (S4 Fig). Prior to incubation, control PU films at Wk0 in normal position (S5 A Fig) and stretched position (S5 B Fig) were completely plain and transparent with no signs of biodegradation. The formation of stains, holes, and deformations increased with time of incubation from Wk2 to Wk4 (S5 Figs). The tensile stress of control PU film was 20.65 Kpa, and that of treatments in Wk1, Wk2, Wk3, and Wk4 were 17.31, 19.01, 16.48, and 10.45 Kpa, respectively, suggesting that tensile strength decreases significantly with exposure to *A. flavus* G10 (S4 B Fig). Differences were highly significant (p=0.005) across weeks. The longitudinal strain for the PU film at Wk0 was 0.8713, and those of Wk1, Wk2, Wk3, and Wk4 films were 0.8502, 0.9306, 0.9364, and 0.9614, respectively. There were significant differences among strain values at different weeks, and the deformation rate increased from W1 to W4 (S4 C Fig) (p=0.007).

### PU degradation by *A. flavus* G10 and formation of MDA

LC/MS showed a peak with an m/z value of 199.1237 ([M+H], m/z: calculated as 199.1157 for C_13_H_14_N_2_) that appeared in both the positive control (MDA) and in PU films exposed to *A. flavus* G10 (Figs 3A-a, c and e). This showed that the degradation of PU film was accompanied by the formation of MDA. It did not appear in the negative control or control PU films (Figs 3A - b and f).

**Fig 3.**
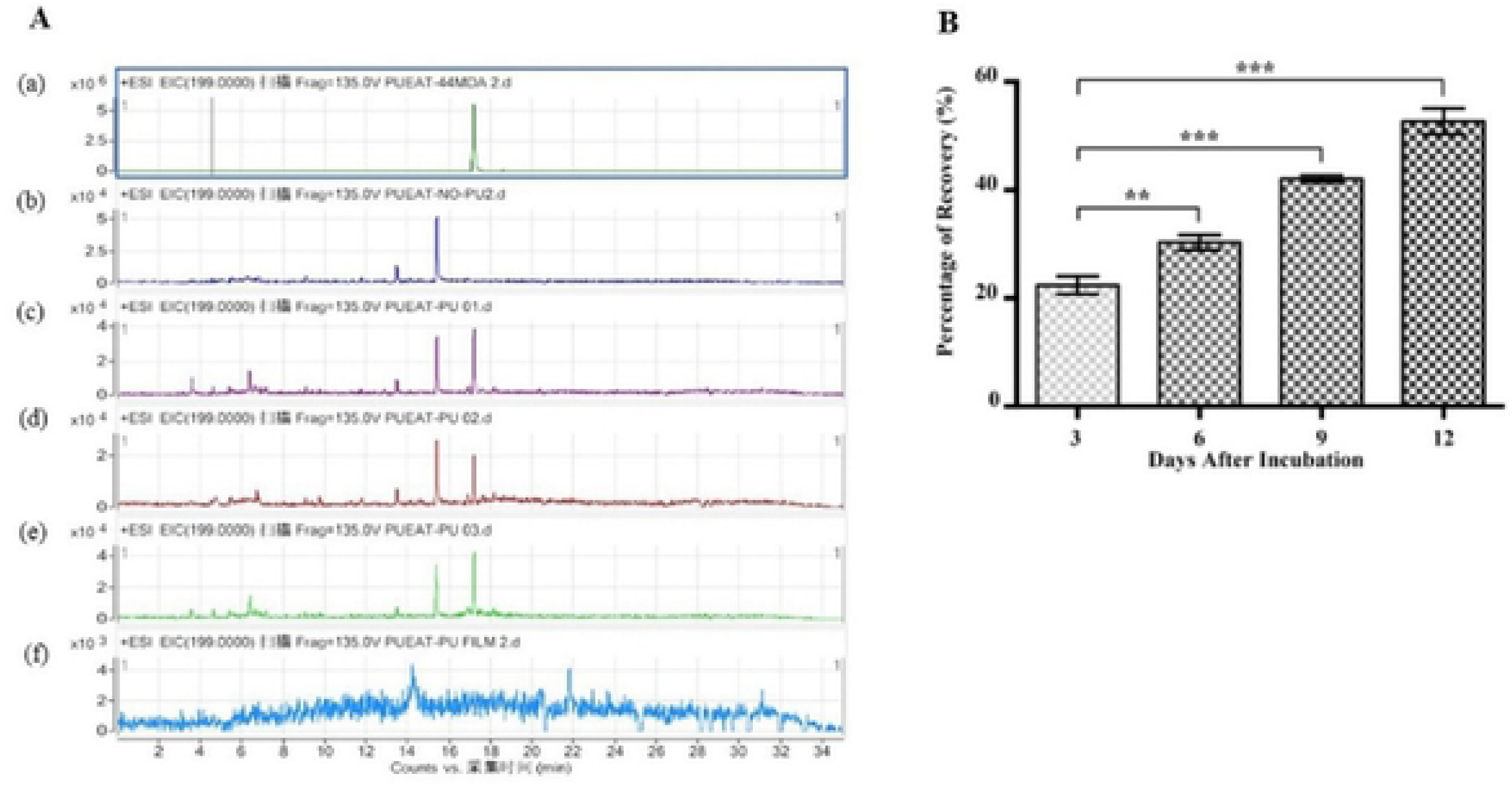
Urethane bond hydrolysis and mineralization of PU films. (A): LC/MS analysis of urethane bond hydrolysis. (a) MDA as a standard compound. (b) MEA medium and fungal mycelium were used as a negative control. (c, d and e) The peaks of the degraded PU films. (f) Represents a non-degraded PU film. (B) Formation of CO_2_ from the PU sheets during their incubation with *A. flavus* G10 in glass jars, for 3, 6, 9 and 12 days. Results from replicate experiments are shown. The data was analyzed using PRISIM 6.0 for significant difference by the One-way-ANOVA (\**P*<0.05, \*\**P*<0.01, \*\*\**P*<0.001).

### Mineralization of PU carbons to CO_2_

Conversion of PU to CO_2_ increased from 22.3% to 52.6% during the 12-day incubation period (Fig 3B). There were significant differences among CO_2_ production across days 3, 6, 9, and 12 (P = 0.01) (Fig 3B).

## Discussion

### Biodegradation of PU by *Aspergillus flavus* G10

This study is the first to report on the isolation and identification of a PU-degrading fungus from the intestine of crickets. *Aspergillus flavus* G10 can degrade PU efficiently, with a mean weight loss of 1.9% per week. The present study offers practical significance for designing and implementing a sustainable PU waste management strategy as well as insight into the usage of biological agents in large-scale biodegradation platforms. Preliminary screening based on gravimetric analysis (data not shown) identified that the biodegradation of PU by *A. flavus* G10 was significantly higher when grown on solid MEA media as compared to the liquid culture media. Analysis of the surface of the PU exposed to *A. flavus* G10 revealed that PU films had been broken down, were replete with numerous holes and cavities, and that fungal spores and hyphae covered the surfaces of these PU films (Figs 1-3). Similar results, such as the formation of holes in the PU film, hyphal growth on its surface, and discoloration, were previously reported for the biodegradation of PU by *A. tubingensis* [11]. AFM also analysis revealed that the surface of the PU films incubated with *A. flavus* G10 had extensive holes, cavities, and pits. Moreover, cavity depth was observed to increase based on lengthier incubation time (S2 Fig). Similar previous work used AFM to analyze surface degradation of PE [32, 17]. These findings provide clear evidence that *A. flavus* G10 can cause significant damage to the physical integrity of PU films.

ATR-FTIR analysis clearly indicated significant shifts in the spectral bands associated with C-H and N-H bonds in the chemical structure of the PU films exposed to *A. flavus* G10. These observations are consistent with past studies that found deformations in the C-H and N-H bonds of PU exposed to *A. tubingensis*, and the observations were attributed to enzymes or radicals released by the fungus [11, 18]. Furthermore, our results demonstrated an increase in the intensity and sharpness of bands assigned to free urethane groups, further confirming PU biodegradation by *A. flavus* G10 (S3 Fig). Significant losses were observed in the mechanical properties of PU films incubated with *A. flavus* G10, indicating a decline in physical integrity of the films as a result of its growth and enzymatic activities (S4 and S5 Figs). Similar results for the loss of tensile strength in PU films due to biodegradation have been previously reported [11, 22, 23, 24, 33]. Yang et al. [32] reported approximately 50% loss in tensile strength due to bacterial biodegradation of polyethylene film. In addition to a breakdown of the mechanical properties of the PU films, those films exposed to *A. flavus* G10 exhibited a steady rate of mass loss over the course of 15 weeks. Past studies have shown similar trends in PE films exposed to bacteria capable of digestion [32, 33, 34], but PU mass loss was higher in this case.

### Optimizing the efficiency of biodegradation

Of the different media we tested, *A. flavus* G10 performed best on MEA, with high biodegradative capabilities. The growth rate of *A. flavus* G10 was three times higher when grown on MEA in comparison with other media used in our study (S6 Fig). However, this preference for MEA by *A. flavus* G10 appears to be quite specific, as it has been shown that *A. tubingensis*, another fungus capable of degrading PU, grew best on MSM media compared to PDA and SDA when isolated from soil [11]. Thus, any trials investigating fungal growth and biodegradative capacity should first test for which media is optimal.

We also showed that temperature, pH, and photoperiod were important factors that influence the performance and biodegradative capacity of *A. flavus* G10. The optimum temperature for *A. flavus* G10 was found to be 25°C, and as temperature increased, the rate of biodegradation began to decline, before becoming totally inhibited at 40°C (S6 D Fig). Khan et al. [11] previously demonstrated high enzyme activities at 37°C in the liquid broth culture of *A. tubingensis*. The difference in the temperature optimum of these fungi may reflect genetic differences. It appears that the ideal pH range for fungal growth is related to the environment from which the fungus was isolated. *Aspergillus flavus* G10 was isolated from an alkaline environment in the gut of a cricket, and was found to perform best in alkaline media, with an optimal pH of 12 (S7 Fig). Confirming our assumption that optimal pH for fungal growth is closely related to the environment from which it originates, the most ideal pH for degradation of PU by *A. tubingensis* was slightly acidic, as it was originally isolated from acidic soil conditions [11]. Numerous photoperiods were tested here and it was determined that light to dark ratios of 8:16 and 14:10 are optimal for the growth and biodegradative abilities of *A. flavus* G10 (S6 Fig).

In addition to optimizing the environmental conditions for *A. flavus* G10, adding salts to the growing media (MEA) was found to have significant effects on the biodegradation process. MgSO_4_ was found to accelerate biodegradation (S7 Fig). Not all salts improved the performance of *A. flavus* G10: CuSO_4_ and ZnSO_4_ were found to inhibit fungal activity. No past studies have investigated the effects of adding salts to fungi degradation, so this research suggests possibilities for future work.

We have shown that the manipulation of media and environmental conditions can significantly influence the biodegradative capacity of *A. flavus* G10; however, we also found that treating the PU films before incubation with *A. flavus* G10 also changed rates and extent of fungal degradation. Past studies have shown that photo, thermal, or chemical treatments of polymers prior to microbial exposure can increase biodegradation [7, 35]. In this study, we performed photo- and chemical-based pre-treatments. We observed that PU films exposed to solutions of either FeSO_4_, 7H_2_O, or NaCl prior to *A. flavus* G10 incubation experienced increased rates biodegradation efficiencies (S7 Fig). Exposure of PU films to certain salts decreases the hydrophobicity of the PU films, thereby enhancing the biodegradation of polymers [7]. Shangguan et al. [36] evaluated the effects of UV on the bacterial biodegradability of bio-polyester poly (3-hydroxybutyrate-co-3-hydroxyhexanoate) and found high rates of biodegradation by exposing polyester powder or film to UV radiation. Consistent with previous research [36], we also found that PU films exposed to UV radiation for 5 min prior to incubation with the fungus showed increased biodegradation (S6 Fig). It has been reported that UV radiation results in surface changes in polymers that microorganisms can access more easily [37]. However, according to this study, a longer exposure time to UV radiation led to a decline in the rates of biodegradation. This may relate to the lower or negligible degradation of PU in landfill and dumping sites.

### Biodegradation of PU by *A. flavus* G10 and production of MDA

Methylene di-aniline (MDA) is known to be the enzymatic biodegradation product of PU [27]. Our results detected high concentrations of MDA in the degradation medium, indicating that *A. flavus* G10 was successful in breaking down PU by secreting hydrolytic enzyme(s). Based on these findings, we suggest that PU biodegradation by *A. flavus* G10 occurs in a two-phase process. In the first phase, the fungus colonizes the surface of the PU and causes physical disruption of PU via hyphal growth. Next, the fungus secretes hydrolyzing enzyme(s) that depolymerize PU into low–molecular-weight compounds. These compounds are utilized by the fungus for its own energy and converted into CO_2_ and H_2_O. In a previous study, the fungal strain PURDK2 was found to be able to hydrolyze both the urethane and urea bonds in the ether-type PU, and utilize the degraded compounds as a carbon source [16]. Our results suggest that production of aromatic amines such as methylene di-aniline (MDA) from PU biodegradation (S3 A Fig) could lead to the chemical recycling of PU; however, further experiments are needed in this area.

### Mineralization of PU

The incubation of PU with *A. flavus* G10 resulted in the production of CO_2_, and after 12 days of incubation, the amount of released CO_2_ was the equivalent of about 52.6% of the C in the PU film (Fig 3). Zumstein et al. [38] have identified that the carbon derived from the biodegradation of poly (butylene adipate-co-terephthalate) polymer is converted into CO_2_ and microbial biomass [38]. They observed the ^13^C-labeled isotopic carbon to develop the synthetic polymer, and the isotopic carbon was seen both in the fungal mycelia and in the evaluated CO_2_. Future studies that build upon on our work using ^13^C-labeled polymer would help us trace the exact fate of PU-derived carbons as well as monitor to what extent the fungus is able to assimilate the C for growth.

## Acknowledgments

This work was supported by the Chinese Academy of Sciences, President’s International Fellowship Initiative (CAS-PIFI), and Key Research Program of Frontier Sciences, CAS. We are thankful to Zhijia Gu, Key Laboratories for Plant Diversity and Biogeography of East China, Kunming Institute of Botany, Chinese Academy of Sciences, for his high level of support in using the scanning electron microscopy for photography.

